# Full quantum mechanical modeling of the enzyme-substrate system: how laccase detoxifies aflatoxin

**DOI:** 10.1101/2020.03.02.973883

**Authors:** Marco Zaccaria, William Dawson, Darius Kish, Massimo Reverberi, Marek Domin, Luca Dellafiora, Takahito Nakajima, Luigi Genovese, Babak Momeni

## Abstract

This work focuses on: 1) the development of a methodology to perform a full Quantum Mechanics (QM) characterization of enzymatic activity; 2) the development of a rational approach to laccase engineering as a food bioremediator. Aflatoxins are among the most dangerous natural carcinogens, and regularly contaminate reserves of staple crops worldwide. Decontamination of aflatoxin-polluted food is of great interest for ensuring food safety, and bioremediation is regarded as the most promising solution. The fungal isoforms of laccase display the rare potential to detoxify aflatoxin by tackling its aromatic moieties.. Yet, because of a generally low efficiency, large-scale application of naturally occurring isoforms has so far been unfeasible. We perform a combination of quantitative experimentation and quantum mechanical modeling on aflatoxin and reveal that: (1) detoxification efficiency is limited by the low enzymatic affinity for the substrate; and (2) aflatoxin is not detoxified by oxidative activity of laccase alone, but requires additional stimulation from the environment. QM modeling also allowed identification of the residues in the laccase tertiary structure that determine affinity of the enzymatic pocket for aflatoxin. We conclude that, for our case-study, a full QM approach is mandatory as a first step towards rational optimization. We detail a feasible approach towards this endeavor and argue that our full QM characterization can serve as a roadmap for enzyme development in other applications pertaining laccase as well as other enzymes.

## Introduction

State-of-the-art characterization of enzymatic-substrate systems often revolves around QM/MM methods that rely on preparatory chemical observations and intuition. QM/MM has shown success in many cases, but its failure goes unnoticed and leaves us with no alternatives for more in depth investigations whenever the relevant dimensional scale exceeds few hundred atoms. Performing a full QM characterization of enzyme-substrate interaction would allow an agnostic approach to enzymatic characterization, independent of the knowledge necessary to hypothesize proper QM regions in the model [1], [2]. Moreover, a full QM approach would allow validation of previous QM/MM models and enable high detail characterizations of less studied/tractable enzyme-substrate systems. The required computational demands for such an approach make it presently unapproachable at the relevant dimensional scale. To attempt a full QM model, a promising candidate would be an experimentally tractable enzyme-substrate system which enables reliable model verification. The present work moves steps in this direction by focusing on the characterization of the laccase-aflatoxin system. The motivation behind this is multi-faceted: 1) Fully mechanistic characterization of the tertiary structure is the mandatory stepping stone to allow a rational approach to the process of Directed Enzyme Evolution [3]. 2) Laccase, an enzymes of great interest in biotechnology [4], is a reasonable choice for developing the know-how of mechanistic enzyme characterization and optimization. 3) In bioremediation, current state-of-the-art is inadequate for targeting aflatoxins—among the most dangerous food pollutants [3].

When fungi grow on food commodities such as corn, cereals, rice, and peanuts, they can produce harmful secondary metabolites called mycotoxins [5]. Among mycotoxins, aflatoxins (AFs) produced by *Aspergillus* species are the most dangerous natural pollutants [6], being associated with several diseases, from liver damage and liver cancer [7] to childhood stunting [8]. Aflatoxin contamination is thus a major food safety concern. To protect consumers, contaminated foodstuff is disposed of every year; decontamination is costly and unsafe in industrialized countries, and often unfeasible in developing ones [9]. In the European Union, the mitigation of mycotoxins levels can be applied on compliant batches, whereas compromised commodities are irredeemable and unsellable Food recovery through bioremediation has been proposed as the most promising long-term alternative to address this concern [10].

One bioremediation strategy is to take advantage of natural occurring enzymes. Among available options, laccase is a versatile and environment-friendly candidate with the potential to detoxify aflatoxin-contaminated food commodities [11]–[13]. Laccase itself has been studied widely before, and is regarded as the “ideal green catalyst” and a pivotal enzyme in biotechnological research [4]. The enzyme is a monomeric multicopper oxidase of relatively small molecular weight, first identified in *Rhus vernicifera* at the end of 19th century in Japan [14]. The enzymatic function consists of a single-electron oxidation of a substrate, coupled with a four-electron reduction of molecular oxygen to water. Laccase is found across several interkingdom taxa: plants, bacteria, archaea, animals, and fungi [15]. The alignments of interkingdom, phylogenetically distant laccases have allowed the identification of the shared signature sequence encoding the four copper ligands of the active site [16]. One of laccase’s defining traits is its versatility: its biological role ranges from catalysis (e.g. lignin degradation by white rot fungi) to polymer synthesis (e.g. chitin fixation in the exoskeleton of Arthropoda). Across natural variants, the fungal isoforms have the highest redox potential (E°)—up to 800 mV [15].

Our goal is to identify the intrinsic limitations of laccase as an aflatoxin detoxifier to devise approaches to overcome them. In this research, particular attention is given to the detoxification of aflatoxin B_1_ (AFB_1_), the most carcinogenic of the AFs. For detoxification experiments, we employed the laccase from the Basidiomycetes *Trametes versicolor* (TV), a fungal species whose ecological niche is tailored around laccase-mediated lignin degradation [17]. We perform an in-depth analysis of the degradation of the main target molecule, AFB_1_, by TV laccase to identify the mechanisms behind reaction bottlenecks. Our data also include experiments on a second AFB_1_ congener, aflatoxin G_2_ (AFG_2_), to compare and contrast our findings.

We first constructed a preliminary phenomenological model based on laccase activity on AFs *in vitro*. The model highlights two salient points: i) laccase’s efficacy against AF is not limited by the redox potential of its active site, rather by poor affinity for AF as a substrate; ii) AFB_1_, unlike AFG_2_, deviates from the established Michaelis-Menten dynamic characteristic of laccase activity [18]. Thus, improving affinity appears to be the best option to achieve successful large-scale application. At the same time, remarkable differences in the degradation of AFB_1_ and AFG_2_ (the latter being 10-fold more efficiently degraded) imply that affinity improvement would not be achieved only by specializing the enzyme towards a general category of compounds (e.g. hydrocarbons, aromatic non phenolic structures, or even aflatoxins as a whole). Instead, it will require a highly-detailed approach that is capable of distinguishing between different congeners.

We proceeded with a quantum mechanical (QM) modeling approach focused on simulating the effects of a laccase-like (single-electron) oxidation of a target aflatoxin molecule. The model is based on density functional theory (DFT) calculations of the neutral and oxidized molecule. In particular, we study the selectivity properties of the gas-phase toxins by analyzing the Fukui function to understand what parts of the toxin molecule might play a role during toxin oxidation. Results show that i) laccase does not achieve aflatoxin degradation through single-electron oxidation only; an ulterior environmental stimulation must happen to achieve detoxification. ii) The observed discrepancies in degradation can at least partially be attributed to the intrinsic properties of the two toxin variants.

Moving forward, we turned to a full ab initio QM model of enzyme-substrate interaction over models that would rely on chemical intuitions. By including laccase in the QM model, we i) identified the amino acid residues pivotal to aflatoxin degradation for each aflatoxin variant; ii) categorized the identified residues based on their degree of enzyme-substrate influence; iii) confirmed that interactions between laccase and the two aflatoxin variants are variant-specific in terms of their strength and localization. We conclude that rational engineering of a laccase-based aflatoxin bioremediator requires a detailed mechanistic approach. We present an effective mechanistic way to inform rational enzyme specialization to target substrates of interest that could contribute to other case studies that require high detail characterization.

## Results

### Laccase is a more effective degrader of AFG_2_ than it is of AFB_1_

The lactone ring in the chemical structure of aflatoxin is the main element responsible for its toxicity [19]. The lactone ring also confers natural fluorescence to the aflatoxin molecule. As a result, aflatoxin concentration and toxicity can be fluorimetrically assayed. In this work, we will define as detoxification a reaction leading to the breakage of the lactone ring in the aromatic structure of AFs. We can thus correlate successful aflatoxin detoxification to loss of native fluorescence. This assay can be used for both AFB_1_ (the most dangerous among aflatoxins and main target of the bioremediation effort) and AFG_2_ (used for comparison).

The fluorimetric assay highlights two different degradation trends for AFB_1_ and AFG_2_ by *T. versicolor* laccase. AFB_1_ fluorescence readout follows a decreasing rate that, after about 10 hours, changes into a slower trend that remains constant for all the experimental time. Overall degradation over 96 hours is about 12% of the original quantity (Fig. 1A). AFG_2_ degradation, in contrast, displays a consistent trend, leading to complete degradation of AFG_2_ after 96 hours (Fig. 1B).

**Figure 1.**
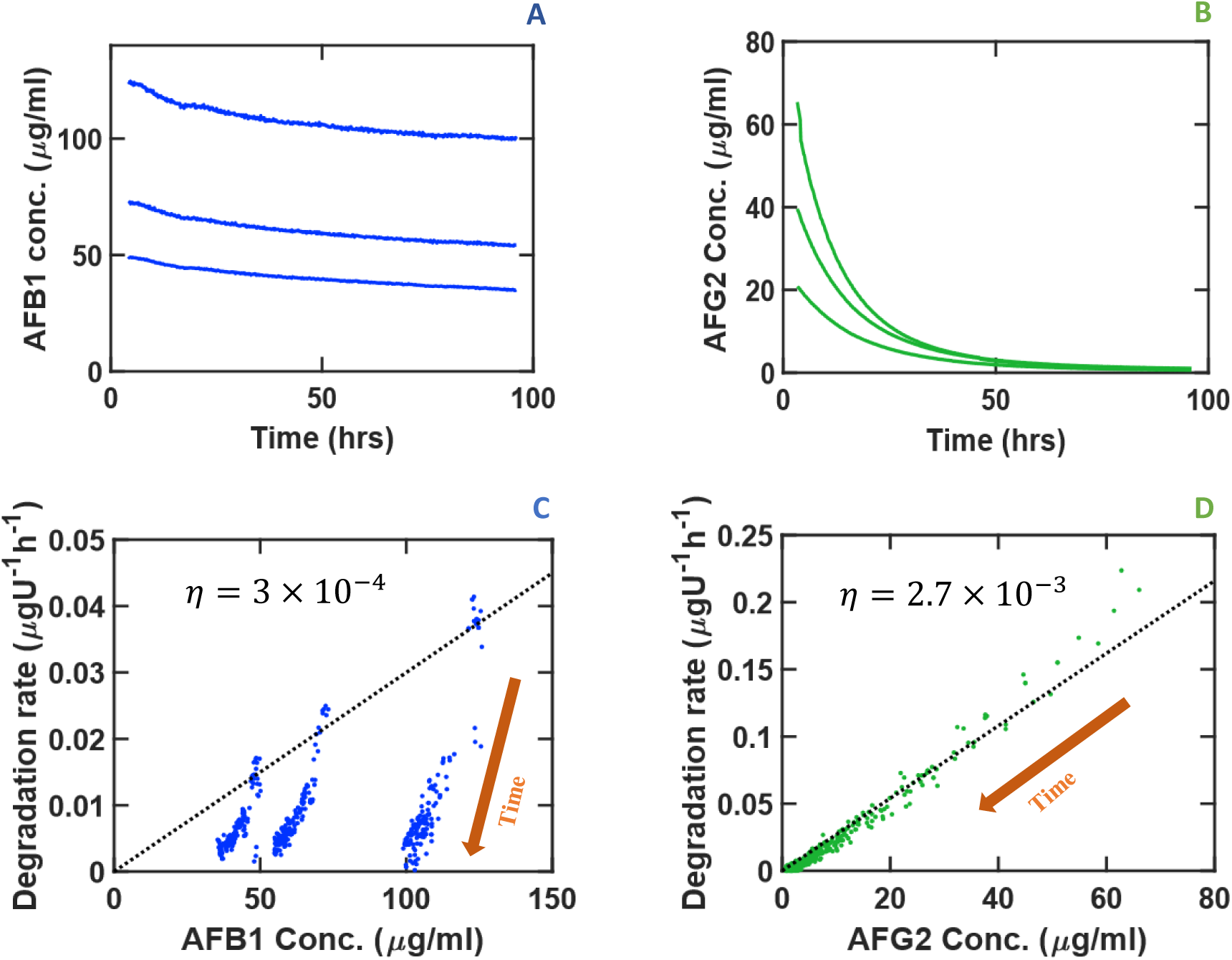
The degradation of (A) AFB_1_ and (B) AFG_2_ by laccase highlights the difference in detoxification efficiencies even between aflatoxins with similar structure. A subset of points from (A) and (B) is randomly selected and represented in (C) and (D) to calculate the local degradation rates. Degradation rate of AFB_1_ (C) is almost an order of magnitude lower than that of and AFG_2_ (D) at comparable concentrations. Dotted lines in (C) and (D) illustrate the prediction of the model. Direction of time is represented in (C) and (D) to highlight the decrease in toxin concentration as a result of degradation.

### Laccase has higher affinity and a higher detoxification rate for AFG_2_ over AFB_1_

In our model, we assume that laccase degrades AFs following the Michaelis-Menten equation:

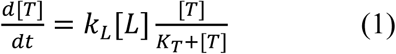

in which [*T*] is the toxin concentration (in *μg*/*mL*), [*L*] is the laccase concentration (in *U*/*mL*), *K*_*T*_ is the Michaelis-Menten constant (in *μg*/*mL*), and *k*_*L*_ is the degradation rate by laccase from enzyme-toxin associated state (in *μg*/*U*/*hr*).

In the limit that the toxin concentration is much lower than the Michaelis-Menten constant (*K*_*T*_ ≫ [*T*]), the equation will be simplified to

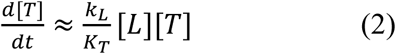

To test if this assumption is valid, from the experimental data we define detoxification efficiency 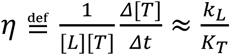 where *Δ*[*T*] is the change in the toxin concentration in a small time-step *Δt*. Since we can measure [*T*] experimentally over time, we can calculate *η. η* appears to be constant throughout degradation, suggesting that *K*_*T*_ ≫ [*T*] is a valid approximation (Fig.1C-D). Calculating the degradation kinetics from Eq (2) and using the value of *η* estimated from experimental data, the algorithm accurately approximates the measured kinetics (Fig. 1C-D), further confirming that this model is suitable for representing aflatoxin degradation by laccase in the case of variant G_2_ throughout the experimental time. However, variant B_1_ adheres to the Michaelis-Menten trend only for a short time before entering a slower, not Michaelis-Menten-like degradation dynamic. Thus, compared to AFG_2_, and other known substrates of laccase [18], AFB_1_ reacts uncharacteristically.

The finding that, at relevant concentrations of the toxin, we get *K*_*T*_ ≫ [*T*] can be interpreted as relatively poor activity by laccase for degrading the toxin. If we consider the association and enzymatic activity in the standard view [20]:

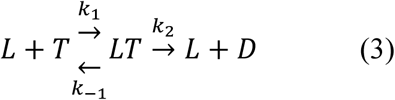

where *D* is the degraded toxin, 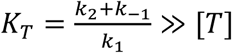 means *k*_2_ + *k*_− 1_≫ *k*_1_ [*T*]. This can be interpreted as low affinity of the enzyme for the aflatoxin, AFB_1_ and AFG_2_ alike, as the rate of association is much smaller than the rates of degradation/dissociation. This low affinity suggests that laccase is naturally not well-adapted to degrade aflatoxin.

To assess the consistency of degradation, we tested 50 U/mL of laccase at increasing intervals of AFB_1_ and AFG_2_ concentrations. Degradation rate appears to be consistent across different concentrations for AFG_2_, in line with our simple model of enzymatic degradation (Fig 1D). For AFB_1_, the model approximates the experimental evidence early on, but the degradation rate drops later (Fig 1C). We make two main observations:

#### 1. Differences in enzyme/substrate affinity between AFB_1_ and AFG_2_

*η* represents the degradation efficiency, which describes how the substrate is affected by the active site and consequential lactone ring opening. For AFB_1_, *η*= 1.2 × 10-5; in the case of AFG_2_, *η*= 1.1 × 10-4. This means that laccase is approximately ten times less efficient at detoxifying AFB_1_ compared to AFG_2_.

#### 2. Differences in degradation dynamics

AFG_2_ degradation rate proceeds uniformly across all the experimental time. Conversely, AFB_1_ degrades with considerably less efficiency after about 10 hours. This suggests a different mode of interaction between the enzyme and the two different toxins.

### QM model suggests possible explanation to the observed differences in affinity and degradation rates between AFB_1_ and AFG_2_

To investigate the difference between AFB_1_ and AFG_2_, we use a QM model (see Methods for details) to analyze these molecules in the gas phase. The model highlights the isosurfaces of the Fukui functions of the two molecules, exposing the sites vulnerable to a hypothetical one-electron oxidation such as the one laccase would perform (Fig. 2). For both molecules, the model reveals that the susceptible areas are neither on the lactone ring nor on its immediate proximities. Thus, the oxidation cannot occur on the lactone ring itself. Nonetheless, the lactone ring opens up after the toxins are exposed to laccase, as can be inferred by the loss of toxin fluorescence and experimentally confirmed by LC-MS analysis of the degradation products (Fig. S2-S3). Oxidation and detoxification therefore do not coincide, although they are interdependent events. The model also reveals that the AFG_2_ molecule displays a more delocalized Fukui function than AFB_1_, and is therefore susceptible to oxidation in a larger number of sites than AFB_1_. This aspect could at least partially explain the differences in the *η* values identified through the previous analysis between AFG_2_ and AFB_1_. In the QM model, in the absence of an environmental stimulation, the lactone ring will not spontaneously open after the toxin is oxidized. Once we add the simplest representative form of an environmental stimulation using a hydrogen atom, the toxin spontaneously undergoes a structural rearrangement that leads to the formation of an epoxide in the terminal ring. If, however, such environmental stimulation is localized in the immediate proximity of the lactone ring, then the structural rearrangement will cause ring-opening (Fig. 3), without the need for the system to overcome a barrier. The aflatoxin G_2_ has a lower free-energy conformation post-ring breakdown (−1.71 eV compared to oxidized state) over aflatoxin B_1_ (−1.34 eV compared to the oxidized state), which could suggest a slightly higher tendency towards this state.

**Figure 2.**
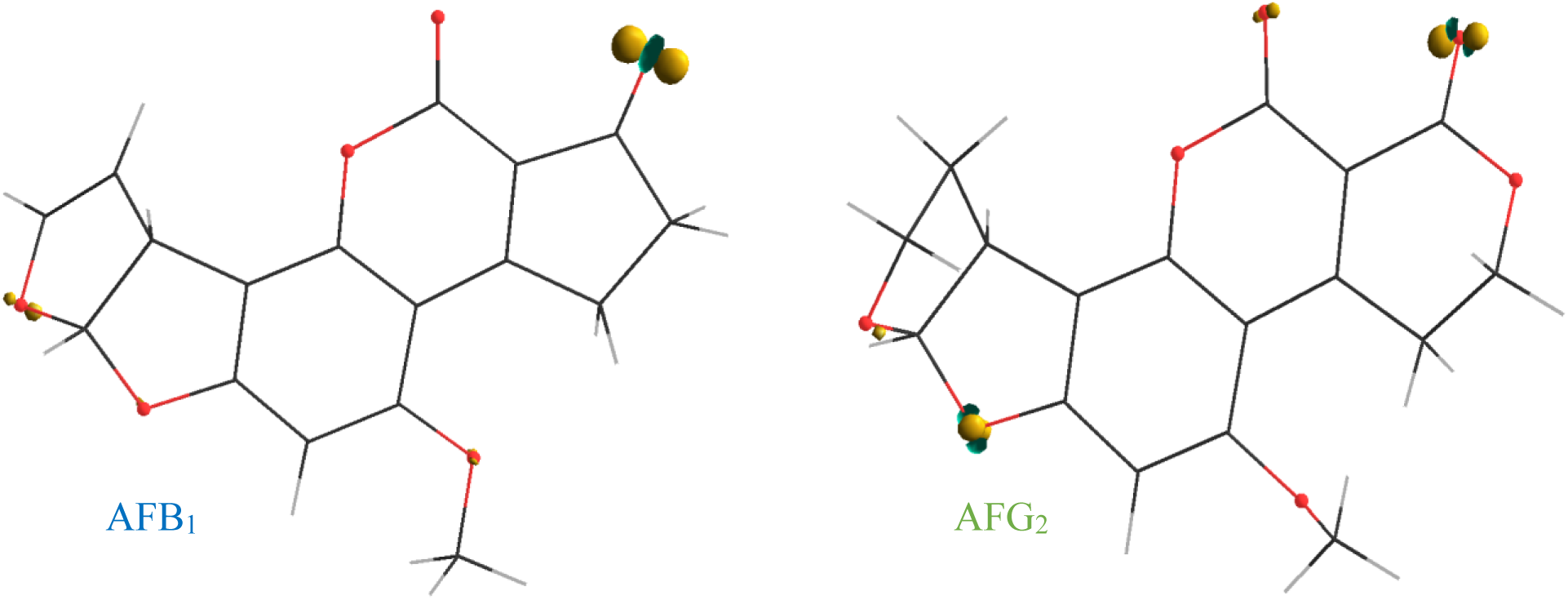
Isosurfaces of the Fukui functions of the gas phase of AFB_1_ and AFG_2_ indicate the sites prone to oxidation. Hydrogen: GRAY; Oxygen: RED; Fukui isosurfaces: YELLOW (+) and BLUE (−). On both molecules, oxidation cannot occur in proximity of the lactone ring. Detoxification (lactone ring opening) is therefore an indirect consequence of oxidation. AFG_2_ molecule has a more delocalized Fukui function, and is spatially more vulnerable to oxidation.

**Figure 3.**
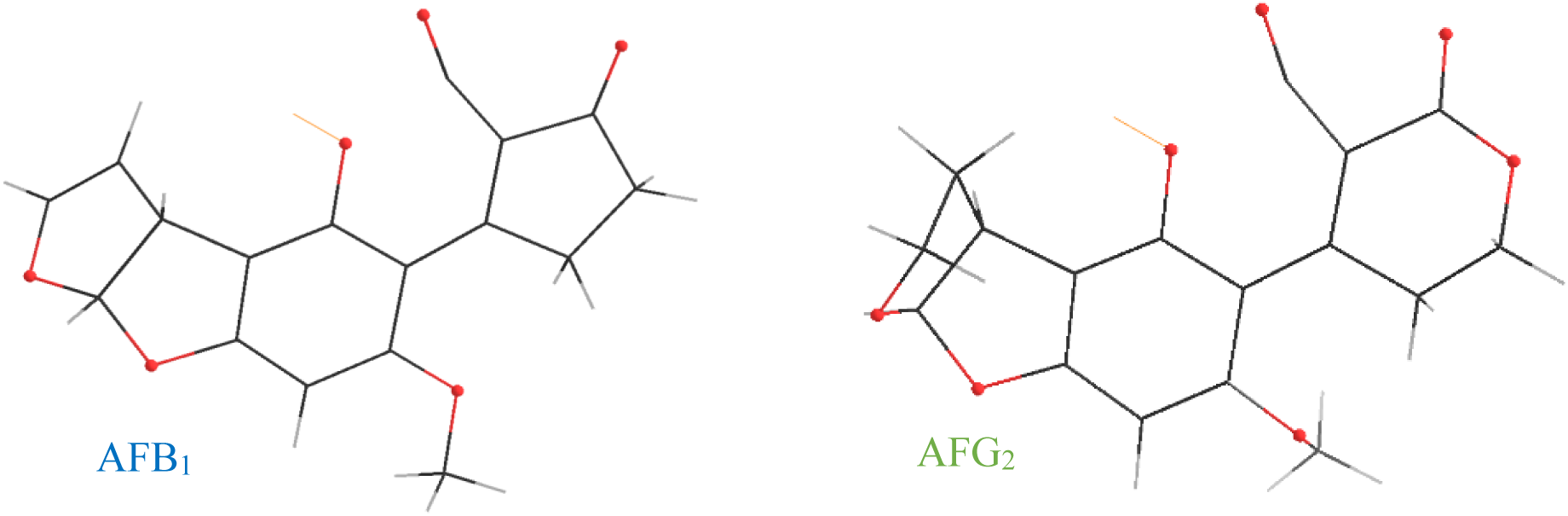
Conformations of the AFB_1_ and AFG_2_ during the simulated lactone-ring opening process indicate the position of environmental stimulation (the attacking hydrogen is shown by the yellow bond). We present a snapshot of the geometry optimization procedure of the oxidized aflatoxin + H system, showing how this would lead to a ring opening.

### Analysis of the binding site shows substrate-specific interaction

To further study the difference in reaction dynamics between the two toxins, we perform modeling of the full docked toxin-enzyme system. Docking positions were taken from a previous study [12], and supplemented with additional poses for the G_2_ toxin (see Methods for details). Among the three isoforms of laccase with different binding sites (i.e. beta, delta and gamma), we found that AFB_1_ interacted favorably with the beta and delta isoform whereas AFG_2_ only interacted favorably with beta. The binding poses of AFB_1_ and AFG_2_ with the beta and delta isoforms are nearly identical with the delta pose being slightly tilted (Fig. 4). The distance between the T1 copper and the shared Fukui function is also similar for each orientation and is oriented outwards from laccase (Tab. 1). However, AFG2 has a second oxidation site which is oriented inward and is significantly closer to the copper. Other recent work [21], [22] has suggested that substrates should dock within 5 angstroms of the histidine residue of the catalytic triad of Laccase (HIS:458 for beta and HIS:479 for delta). Similar to the copper distances, it is only the unique Fukui function site of AFG_2_ that is in a suitable position. Thus, while geometric considerations may be sufficient to improve affinity for these two similar substrates, the activity cannot be understood without considering the electronic interactions.

**Table 1.**
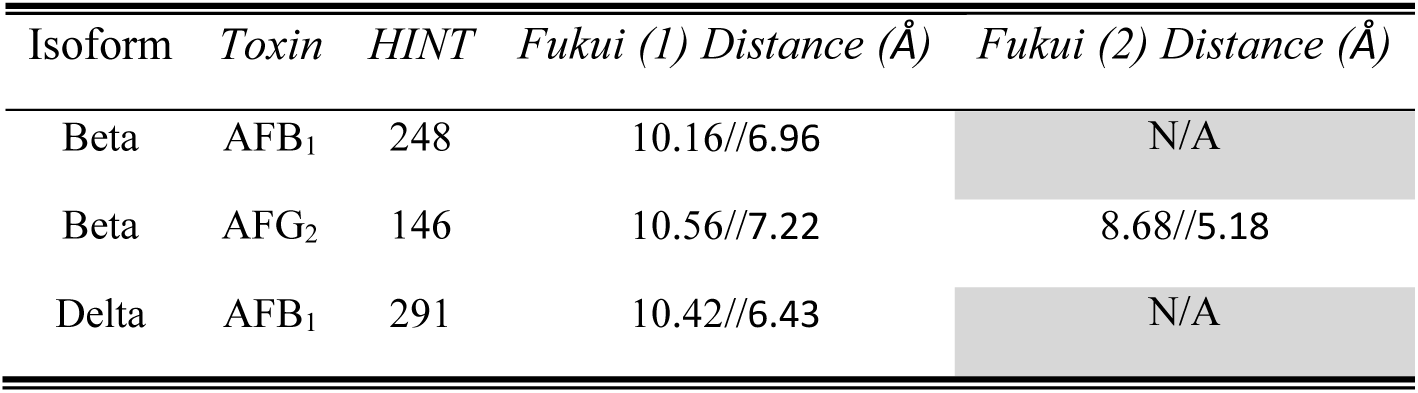
Binding positions of the two toxins with different isoforms of laccase. The HINT docking score provides a measure of affinity (see Methods). We also report the distance between the copper atom//HIS residue from the catalytic triad and the Fukui function between the toxins (1) and the unique Fukui function on AFG_2_ (2).

| Isoform | Toxin | HINT | Fukui (1) Distance (Å) | Fukui (2) Distance (Å) |
| --- | --- | --- | --- | --- |
| Beta | AFB<sub>1</sub> | 248 | 10.16//6.96 | N/A |
| Beta | AFG<sub>2</sub> | 146 | 10.56//7.22 | 8.68//5.18 |
| Delta | AFB<sub>1</sub> | 291 | 10.42//6.43 | N/A |

**Figure 4.**
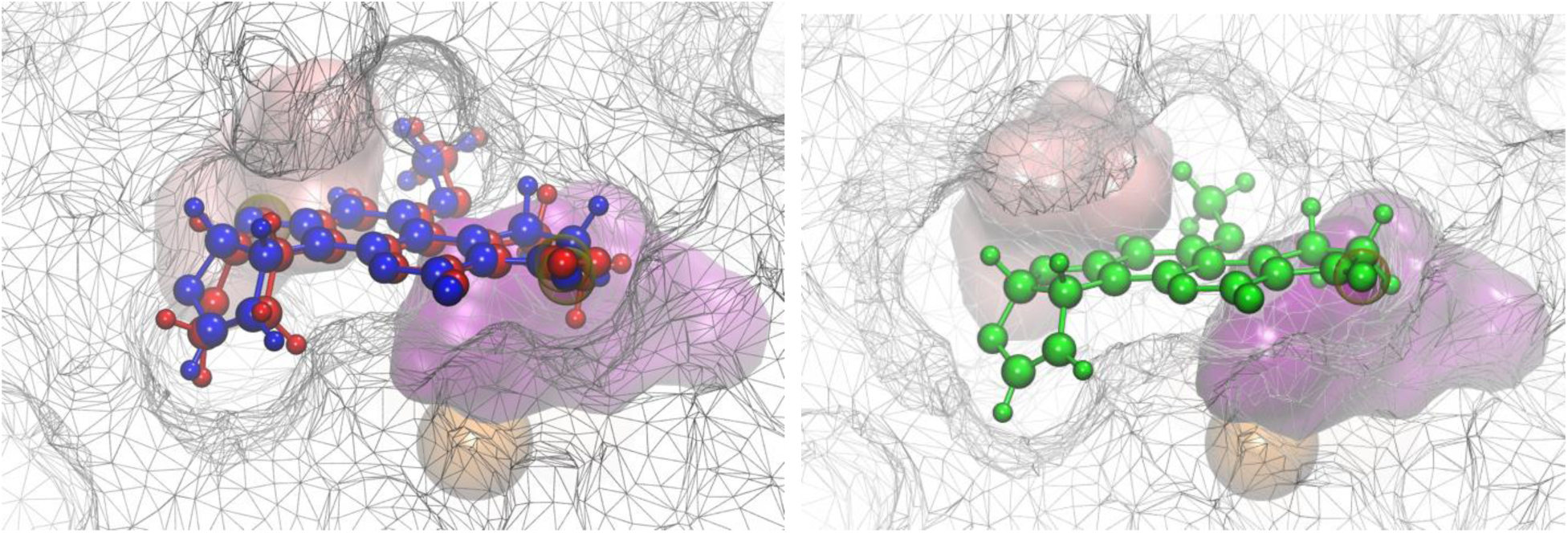
Docking of AF poses. The beta isomers are plotted on the left with AFB_1_ in blue and AFG_2_ in red. The delta isomer with AFB_1_ in green is plotted on the right. The protein is represented by a wired isosurface and the coppers are represented as orange balls. The HIS and ASN amino acids of the catalytic triad are highlighted as solid pink and purple surfaces respectively. The atoms on which the Fukui function reside are highlighted in yellow.

By performing QM modeling of the full system, and post-processing using the Fragment Bond Order tool described in our earlier publication [23], [24] we can generate a more detailed, graph-like view of the electronic interactions with laccase site (Fig. 5). These interactions can be used to define a substrate specific binding site. In this framework, we observe differences in the interaction strength of the toxin and copper depending on the particular configuration. We caution that a more realistic set of atomic positions than those presented here would be required to fully understand this effect. The substrate binding site definitions presented here can guide such future studies.

**Figure 5.**
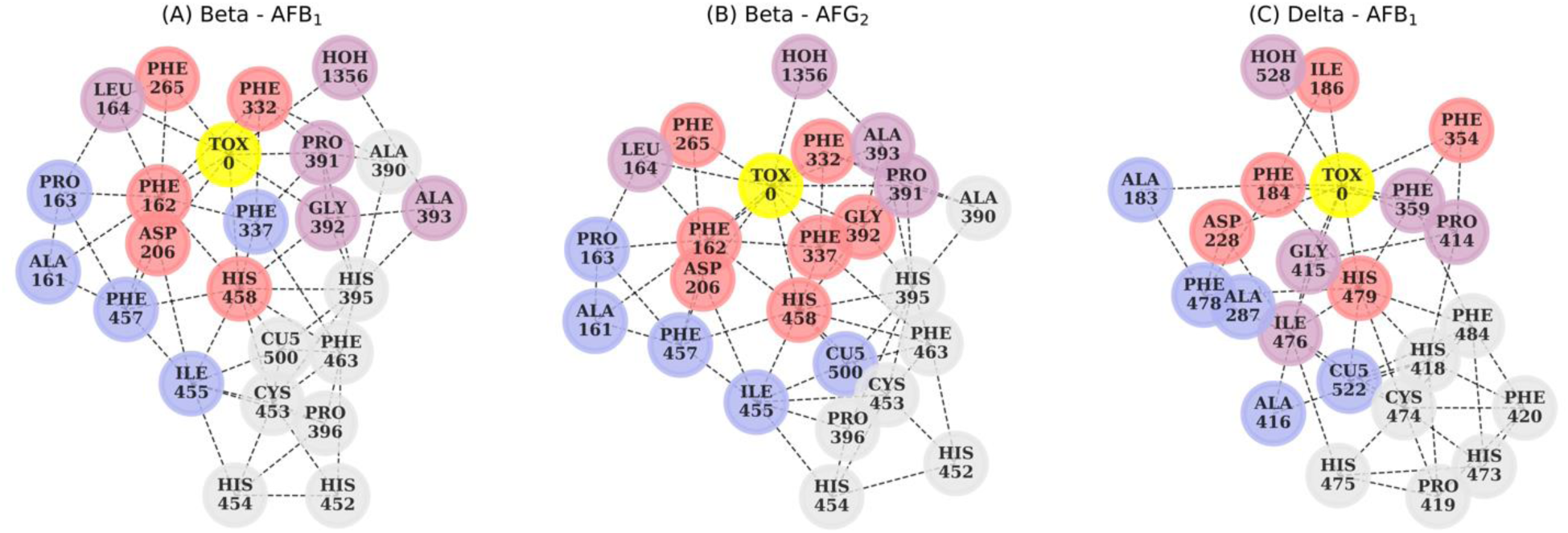
Graph representation of the substrate specific binding sites. The toxin is labeled “TOX 0” in yellow, and the lone copper “CU5 522”. Nodes are labeled according to the residues in the PDB files (Supplementary). The strength of the interaction with the toxin is colored with red as the strongest, followed by purple, blue, and finally gray (weakest).

## Discussion

Aflatoxin contamination is a major concern among food-safety issues. AFB_1_ is classified as a Group 1 carcinogen by the International Agency for the Research on Cancer (IARC), which makes it one of the most dangerous natural compounds [25]. AFs are ubiquitous contaminants of staple crops and their presence on field is regarded as unavoidable [26]. Developing countries in particular face enormous health risks to this day [9]. In addition, climate change towards higher temperatures increases the risk even in areas where aflatoxin contamination has so far been scarce [26]. Therefore, the advantages of employing a well characterized, environment-friendly enzyme like laccase for detoxification have long been stated, and feasibility concerns have concurrently been raised. Once laccase efficacy at degrading aflatoxins had been proved *in vitro* [11], efforts were directed at enabling application. The laccase we employ (Sigma-Aldrich), is isolated from *Trametes versicolor* cultures. *T. versicolor* (or turkey tail, in layman terms) is a Basidiomycetes that grows on rotting wood; it employs laccase to degrade lignin to adhere to the wood and scavenge it for nutrients. Therefore, laccase activity is central to its ecological role, and it is reasonable to assume that it has undergone a constant selective pressure to refine its interaction with lignin, an interaction often mediated by accessory interactors [27]. The progressive evolution of a lignin degrading, laccase-based enzymatic system has led to an active site of the highest redox potential among multi-copper oxidases, which is a good asset to degrade the aromatic moieties of aflatoxins. However, as our data highlight, this adaptation, not surprisingly, disregarded high affinity towards AFB_1_. If an artificial selection regime was attempted to direct laccase towards AF affinity, the very centrality of laccase in the metabolism of *T. versicolor* (and other lignin-degraders) could pose a challenge. Therefore, the most effective works in this context have so far relied on enhancing laccase synthesis through isolating the best natural producers and exposing them to laccase-inducing culture conditions [11]. Culture filtrates would then be collected, concentrated in their laccase-rich fraction and tested for application as an anti-aflatoxin treatment within the food production chain. The methodology behind the previous works is as simple as it is sound: identify a cultivable producer of a laccase of the highest redox power among the natural isoforms; enhance its laccase production regime and isolate culture filtrates at extreme laccase concentration. However, to this day, research is not at the point of streamlining a feasible, cost-efficient solution through such an approach.

The present work provides its contribution by addressing the issue from a novel angle, which is mechanistic by vocation and interdisciplinary by necessity: a bottom-up understanding of the detoxification process. We started by identifying the relevant variables in the dynamics of the laccase/aflatoxin interaction. In our experiments, a simple model of enzymatic degradation highlighted how enzyme abundance, laccase oxidative power, and substrate affinity were sufficient to describe the dynamics of degradation. Moreover, the model suggested that affinity and degradation rate bear the same weight in providing the function of interest: i.e., when contributing to the overall degradation rate, an increase of *n*% in one variable equates to an *n*% increase in the other. Importantly, a comparison with a different aflatoxin variant exposed substrate affinity as the limiting factor in the detoxification potential of the natural enzyme isoforms. In this regard, it is interesting to note that the laccase degradation of the AFG_2_, while much faster than AFB_1_, is still lacking the efficacy necessary to warrant realistic applications, with affinity still constituting the real drawback.

For bioremediation to have a realistic chance, it has to be seamlessly implemented during the current food production process without disrupting it. With this in mind, the best context for laccase-mediated aflatoxin bioremediation would be during the water washing-step of food commodities. For that, AFB_1_ detoxification would need to be achieved in no longer than about 3 hours, in a slightly acidic (pH of 6.5), aerobic, liquid environment at room temperature. At the same time, QM models of laccase active site and surrounding regions aimed at informing rational enzyme engineering have so far improved a function of interest by about 3-fold at best [28], [29]. Our data to this point expose one main issue: natural affinity for AFB_1_ is so low that a 3-fold increase would still make for a subpar bioremediator. Thus, in virtue of the equivalent weight of the variables in the model, evolving a laccase producer three times more efficient than the WT, while experimentally remarkable, would still leave us far from enabling large-scale application. Moreover, the degradation rate by *T. versicolor* laccase is about 10 times higher for AFG_2_ over AFB_1_. This difference is remarkable and surprising, if we consider how structurally similar the two substrates are. These observations suggest that, to optimize laccase degradation of AFB_1_, a study to the highest level of detail would not be an affectation, but a necessity. This necessity can be addressed through a mechanistic, quantum mechanics-based approach.

Our QM description provides three conclusions: 1) The lactone ring cannot open unless oxidation happens first. 2) Additional environmental stimulation with the lactone ring of the oxidized toxin is required to cause ring opening: i.e. laccase does not directly interact with the lactone ring to achieve detoxification and does not need to, because oxidation is achieved elsewhere. 3) Laccase’s higher affinity for AFG_2_ over AFB_1_ can be partially explained by the more delocalized Fukui function of AFG_2_, which makes AFG_2_ prone to oxidation from more than one side. This last point suggests that aflatoxins degradation potential of natural laccase may be limited, because it depends on the toxin’s intrinsic traits. Nonetheless, before ruling out the application of laccase for aflatoxin degradation, the mechanistic details of the enzyme-substrate compound dynamics need to be formalized.

QM modeling is by definition a mechanistic approach and can thus be used to validate any alternative empirical approach. Reservations about its employment are only limited to its feasibility: present-day computational power cannot reliably model QM-described atomic interactions of more than a few hundred atoms. It is evident that gas-phase QM calculations of the neutral and oxidized toxin may only provide speculative results of the actual processes eventually leading to detoxification. Recently, it has been shown that with a sufficient set of descriptors from docking, and quantum mechanical modeling of the gas phase substrate, it is possible to predict laccase affinity on a wide class of systems [22]. The specific case shown here, however, demonstrates a need for the combined modeling of the enzyme and substrate in order to properly predict degradation. In this view, computational protocols like the quantum mechanics/molecular mechanics (QM/MM) techniques (as employed for laccase in [28], [29]) may be employed. However, such techniques are based on chemical intuition for preliminary identification of the active site region which may be altered and modified by the actual conformation of the enzyme-substrate complex, as we have demonstrated in this paper for the particular case of AFB_1_ and AFG_2_.

It is thus appealing to consider a full QM approach which would shed light on various relevant questions, for instance: 1) Is the oxidation rate of the toxin dependent on how the latter approaches the active site? 2) Is the enzyme active site altered by the toxin presence? 3) How accurate are the QM/MM approximations with respect to an unbiased, full QM calculations? In order to provide answers to such questions, it is greatly beneficial to employ a QM approach that is, potentially, able to treat systems up to many thousands of atoms at the QM level of theory. The BigDFT code, employed in this paper for QM modeling, has been proven to be able to tackle KS-DFT calculations of systems up to few tens of thousands atoms [30], [31], and can provide reliable information on the identification of the systems’ fragments and associated physical observables [24]. Work is ongoing in the direction of the full QM characterization of prototypical enzyme-substrate complex conformations.

In the specific case of AFB_1_ degradation, we have presented evidence that selecting for general categories of substrates would likely not be enough, as suggested by the striking difference between the degradation rates of AFB_1_ and AFG_2_. The QM model of the laccase-aflatoxin complex can inform rational selection strategies for efficient directed enzyme evolution. Our analysis has provided a map of the residues that play a direct role in the affinity of the toxin molecule for the enzymatic pocket, and also allowed an estimate of their intensity of interaction. These observations can guide rational engineering efforts by providing insights about the ideal enzyme structure. In the future, the data acquired will allow us an *in silico* generation of all the theoretical structures of laccase capable of degrading aflatoxin to the best possible rate, and also highlight laccase’s intrinsic limitations. The theoretical cues could then be evaluated experimentally by generating the respective isoforms through gene editing. We provide a methodology for a full QM approach that can benefit several case-studies that focus on the mechanisms of enzyme-substrate interaction (e.g. specific instances of antibiotic resistance). Fungal enzymes such as *T. versicolor* laccase are the most promising category of degraders for pollutants of synthetic and natural origin [32]. A successful, rational approach to laccase engineering is, nowadays, not beyond feasibility.

## Materials and Methods

### Fluorescence-based assay of laccase-mediated Aflatoxin B_1_ and Aflatoxin G_2_ degradation

Laccase from *Trametes versicolor* (Sigma-Aldrich CAS80498) was dissolved in acetate buffer (pH 6.5) at a final concentration of 25U/mL. Aflatoxin B_1_ and Aflatoxin G_2_ (Cayman Chemicals) were dissolved in LS-MS grade methanol (Sigma-Aldrich) at 4 different concentration intervals: 3, 30, 50, 100µg/mL. Buffer solutions of laccase and aflatoxins were incubated at 28°C over 96 hours under fast continuous shaking regime in a Synergy™ Mx Multi-Mode Microplate Reader (Biotek), each condition was performed in triplicate. Due to their natural fluorescence, aflatoxin concentration was fluorimetrically assayed (ex 380nm - em 440nm, gain 65 and 50); readouts were acquired every 10 minutes, totaling 577 by the end of the experiment. Controls were used assaying laccase fluorescence in the buffer in the absence of aflatoxins, and AFB_1_ and AFG_2_ fluorescence in the absence of laccase. To convert the fluorescence readout to the corresponding toxin concentration, a calibration curve was used based on measurements at a set of known toxin concentrations (Fig. S1).

### Mathematical modeling of laccase/aflatoxins interactions

The reaction kinetics were simulated using Matlab. The source codes are available on GitHub at: github.com/bmomeni/laccase-aflatoxins-reaction-kinetics

### Data analysis

The method of least squares was employed to fit the Michaelis-Menten kinetics to the experimental data. Degradation rates were estimated by fitting a line to data from the first 100 minutes of the experiments for each initial toxin concentration.

### Docking of AFs with Laccase

The 3D model of *Trametes versicolor* beta isoform laccase was derived from the crystallographic structure of this enzyme recorded in the Protein DataBank having access code 1KYA [33]. The delta and gamma isoforms were derived instead via homology modeling using the beta isoform as a template as previously described [12]. The 3D structure of the aflatoxins were downloaded from PubChem (https://pubchem.ncbi.nlm.nih.gov/) [34], [35]. Before docking analysis, the consistency of atoms and bonds type for proteins and ligand were checked using the software Sybyl, version 8.1 (Certara USA, Princeton, NJ, USA), in agreement with a previous [12] study (REF). Docking simulations were performed using the software GOLD (Genetic Optimization for Ligand and Protein) and each docking pose was rescored using the HINT scoring function [36] (REF) for a better evaluation of the protein-ligand interaction, as previously reported [12].

### Quantum Mechanics-based modeling of a single electron oxidation effect on AFs molecules

QM calculation were performed within the framework of Kohn-Sham Density Functional Theory (KS-DFT) [37], employing the Perdew–Burke-Ernzerhof (PBE) [38] exchange and correlation level of theory. The numerical results were extracted employing the BigDFT code [39], which uses Daubechies wavelets to express the KS orbitals. Hartwigsen-Goedecker-Hutter (HGH) pseudopotentials [40] were used to remove the core electrons orbitals. The use of wavelet basis sets enables one to control the precision of the results within a systematic approach, and at the same time to explicitly consider calculations of charged systems, as isolated boundary conditions are explicitly included in the calculations, without supercell aliasing effects, using the Poisson Solver of the code [41]. A wavelet grid spacing of 0.37 atomic units has been employed for the calculations presented in this work.

The code was used to calculate charged Delta-SCF, and the Fukui functions (FF) are defined as the difference between the neutral ground state electronic density and the corresponding quantity in the (vertical) cationic state.

To calculate the binding site, we performed KS-DFT calculations on the docked enzyme-toxin system using the linear scaling mode of the BigDFT code [27], [28] with the PBE approximation, HGH pseudopotentials, and a grid spacing of 0.4 atomic units. The charge of the enzyme was determined by minimizing the energy of the gas phase enzyme with respect to the number of electrons. Interaction strengths between system fragments were determined using the Fragment Bond Order tool as described in our previous study [23]. The binding site of each enzyme-toxin-pose configuration were determined by including all fragments such that the sum of the fragment bond order of all excluded fragments fell below a cutoff 0.001. The binding site was supplemented by fragments by performing the same analysis on the lone copper. Fragments in red are those that were included when the cutoff of 0.1 was used. Purple and blue used cutoffs of 0.01, and 0.001 respectively, and the gray fragments are the remainder. Edges were drawn between fragments based on a cutoff of 0.01.

## Acknowledgements

BM and MZ were supported by a start-up fund from Boston College and by an Award for Biomedical Excellence from the Smith Family Foundation. This work was supported by the Next-Generation Supercomputer project (the K computer) and the FLAGSHIP2020 project (Supercomputer Fugaku) within the priority study5 (Development of new fundamental technologies for high-efficiency energy creation, conversion/storage and use) from the Ministry of Education, Culture, Sports, Science and Technology (MEXT) of Japan. Experiments presented in this paper were carried out using the Grid’5000 testbed, supported by a scientific interest group hosted by Inria and including CNRS, RENATER and several Universities as well as other organizations (see https://www.grid5000.fr). Additional calculations were performed using the Hokusai supercomputer system at RIKEN. This work was partly supported by Cabinet Office, Government of Japan, Cross-ministerial Strategic Innovation Promotion Program (SIP), “Technologies for Smart Bio-industry and Agriculture” (funding agency: Bio-oriented Technology Research Advancement Institution, NARO)

## Supplementary Information for

### 1. Fluorescence based calibration curve for aflatoxin concentration

Fluorescence based experiments were run as described in the methods section.

**Figure S1.**
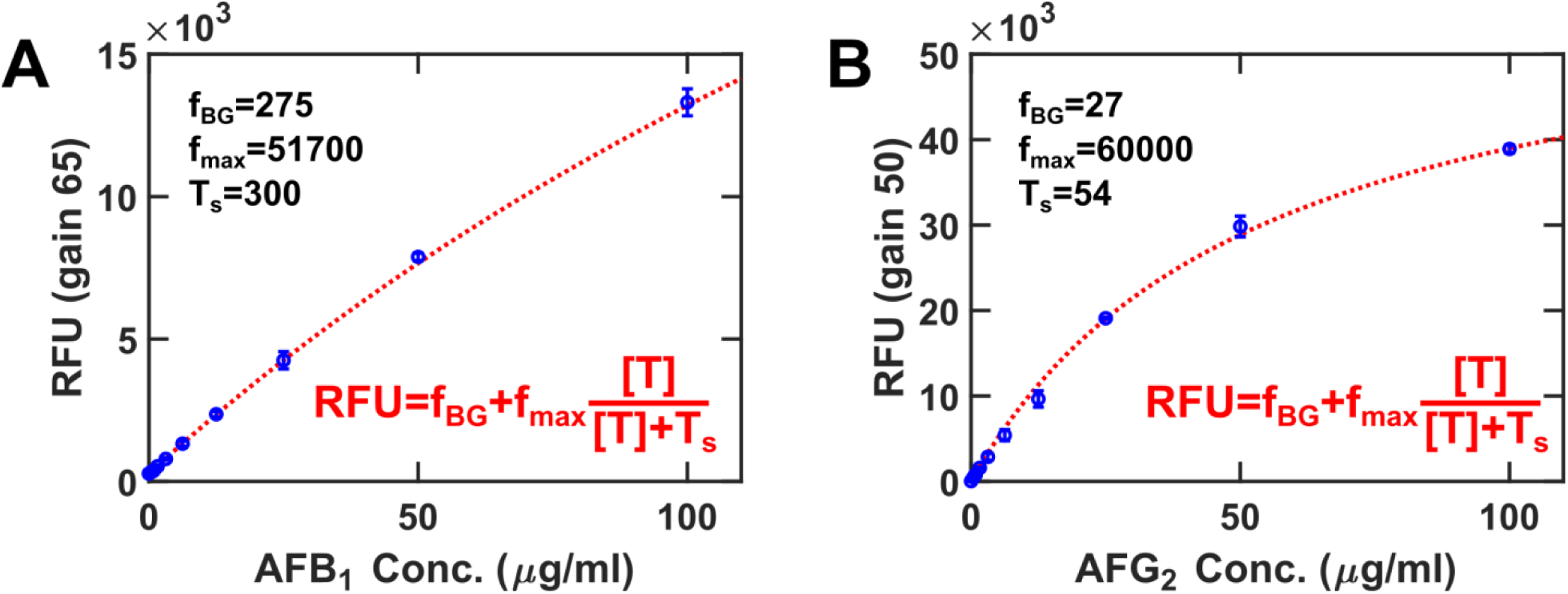
Calibration curves for fluorescence-assayed AFB_1_ (A) and AFG_2_ (B). T represents toxin concentration; RFU represents Relative Fluorescence Units; T_s_, f_BG_ and f_max_ are fitting parameters; the fitting method is Matlab’s Least square.

### 2. Identification of aflatoxin degradation products of laccase activity

#### LC/MS conditions

Column: Kinetex 2.6µm EVO C18; 100 × 2.1mm.

Mobile phase A: Water 5mM Ammonium Acetate, 0.5% Acetic Acid

Mobile phase B: Methanol 5mM Ammonium Acetate, 0.5% Acetic Acid

Flow rate: 350µl/min

UV Wavelength: 354, 360nm

| **Time (min)** | **%A** | **%B** |
| --- | --- | --- |
| **Initial** | **90** | **10** |
| **3** | **90** | **10** |
| **10** | **30** | **70** |
| **10.1** | **10** | **90** |
| **12** | **10** | **90** |
| **12.1** | **90** | **10** |
| **15** | **90** | **10** |

The eluent from the column was directed into the electrospray source of an Agilent 6220 TOF mass spectrometer operated in positive ionization mode. Data was converted into mzML file format and analyzed using MZMine software.

**Figure S2.**
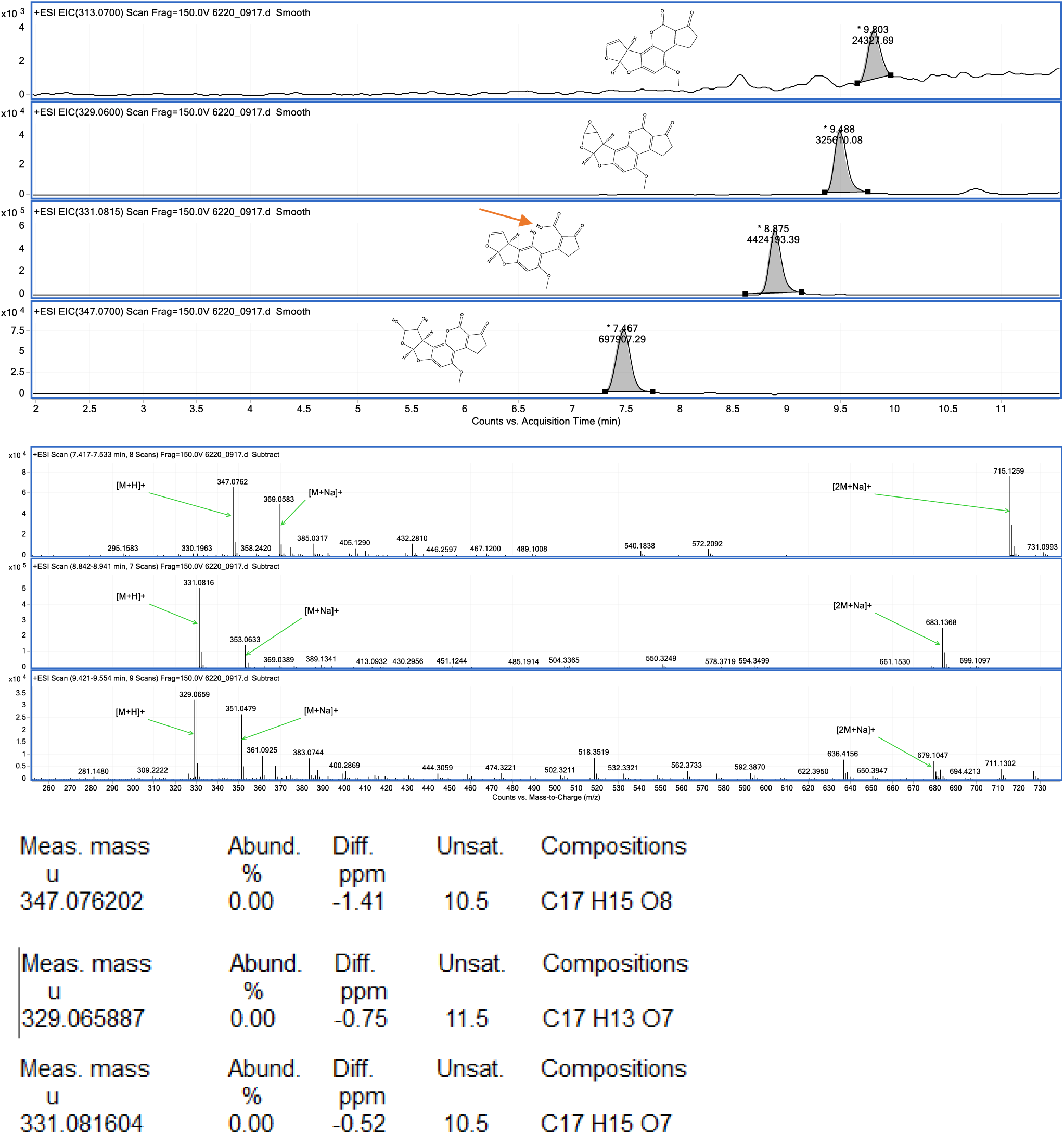
LC-MS analysis of Laccase-mediated AFB_1_ degradation. The fully detoxified molecule is the one with an open lactone ring, highlighted by the red arrow.

**Figure S3.**
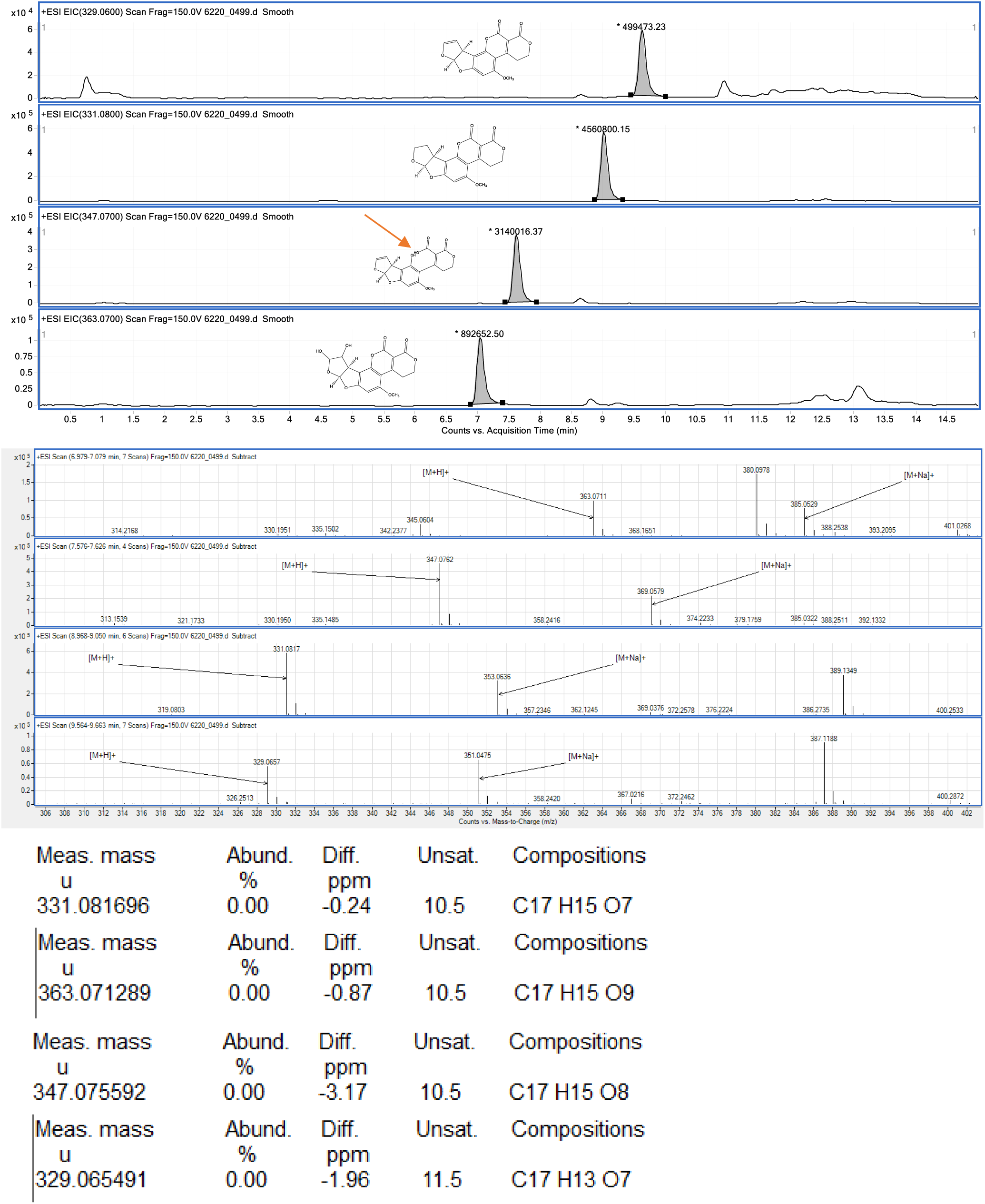
LC-MS analysis of Laccase-mediated AFG_2_ degradation. The fully detoxified molecule is the one with an open lactone ring, highlighted by the red arrow.

